# Accuracy of dipole source reconstruction in the 3-layer BEM model against the 5-layer BEM-FMM model

**DOI:** 10.1101/2024.05.17.594750

**Authors:** Guillermo Nuñez Ponasso, Ryan C. McSweeney, William A. Wartman, Peiyao Lai, Jens Haueisen, Burkhard Maess, Thomas R. Knösche, Konstantin Weise, Gregory M. Noetscher, Tommi Raij, Sergey N. Makaroff

**Affiliations:** Dept. of Electrical and Computer Engineering, Worcester Polytechnic Institute, Worcester, MA, USA; Technische Universität Ilmenau, Ilmenau, Germany; Max Plank Insititute for Human Cognitive and Brain Sciences, Leipzig, Germany; Athinoula A. Martinos Center for Biomedical Imaging, Massachusetts General Hospital, Harvard Medical School, Boston, MA, USA

**Keywords:** electroencephalography (EEG), EEG source analysis, EEG dipole reconstruction, head modeling, boundary element method (BEM), fast multipole method (FMM), adaptative mesh refinement (AMR)

## Abstract

**Objective:** To compare cortical dipole fitting spatial accuracy between the widely used yet highly simplified 3-layer and modern more realistic 5-layer BEM-FMM models with and without *adaptive mesh refinement* (AMR) methods.

**Methods:** We generate simulated noiseless 256-channel EEG data from 5-layer (7-compartment) meshes of 15 subjects from the Connectome Young Adult dataset. For each subject, we test four dipole positions, three sets of conductivity values, and two types of head segmentation. We use the *boundary element method* (BEM) with *fast multipole method* (FMM) acceleration, with or without (AMR), for forward modeling. Dipole fitting is carried out with the FieldTrip MATLAB toolbox.

**Results:** The average position error (across all tested dipoles, subjects, and models) is ∼4 mm, with a standard deviation of ∼2 mm. The orientation error is ∼20° on average, with a standard deviation of ∼15°. Without AMR, the numerical inaccuracies produce a larger disagreement between the 3- and 5-layer models, with an average position error of ∼8 mm (6 mm standard deviation), and an orientation error of 28° (28° standard deviation).

**Conclusions:** The low-resolution 3-layer models provide excellent accuracy in dipole localization. On the other hand, dipole orientation is retrieved less accurately. Therefore, certain applications may require more realistic models for practical source reconstruction. AMR is a critical component for improving the accuracy of forward EEG computations using a high-resolution 5-layer volume conduction model.

**Significance:** Improving EEG source reconstruction accuracy is important for several clinical applications, including epilepsy and other seizure-inducing conditions.

## I. Introduction

EEG source reconstruction, or source localization, consists of locating the neural activity within the brain from EEG-recorded measurements (Knösche and Haueisen 2022). Many excellent open-source software packages implement EEG source localization, including Brainstorm (Tadel et al. 2011), FieldTrip (Oostenveld et al. 2011), MNE (Gramfort et al. 2014), and EEGLab (Delorme and Makeig 2004).

All four packages include BEM-based dipole source localization, typically using three layers extracted from the subject’s MRI: scalp, outer skull, and inner skull. For most practical applications, the resolution of these layers is limited to less than 10 000 triangles per layer. One of the main reasons for this limitation is that these packages use the classical potential-based formulation of the BEM (Geselowitz 1967, Kybic et al. 2005, Ponasso 2023), which involves dense system matrices and as such is unable to compute lead fields of large high-resolution models. Complicating this issue is the fact that the intracortical compartments typically involve very narrow gaps between layers, which leads to numerical inaccuracies that typically need to be resolved with refinement techniques such as *adaptive mesh refinement* (AMR), which is also known as *h*-refinement (Wartman et al. 2024; Weise et al. 2022), or the less common *p*-refinement (Partheymüller, Białecki, and Kuhn 1994), which consists of adaptively increasing the polynomial order of local approximations of the variable of interest (potential or charge) on the mesh triangles.

Yet, current automated human-head segmentation tools (FreeSurfer, Fischl 2012; SPM12, Nielsen et al. 2018) can provide us with much higher resolution skin, skull, cerebrospinal fluid (CSF), grey matter (GM), and white matter (WM) layers, among other tissues. Modern charge-based BEM techniques with FMM acceleration, see for example (Makarov et al. 2018), can compute forward solutions for high-resolution models involving tens of millions of facets, therefore removing the ∼10 000 triangle limitation of conventional BEM. How would the source reconstruction change if we assume the high-resolution models are ground truth? We will investigate this for the canonical EEG problem — single dipole fitting.

There are many studies considering the influence of different uncertainty factors on EEG source localization: head model complexity (Vorwerk, Cho, et al. 2014), conductivity uncertainty (Vorwerk, Aydin, et al. 2019), and white matter anisotropy (Güllmar, Haueisen, and Reichenbach 2010; Bangera et al. 2010) among others. In all these studies, the *finite element method* (FEM) was used for forward modeling, i.e. for generating electrode voltages given a dipole source in the brain. Here, we instead use a high-resolution BEM model, coupled with FMM acceleration, to compute the forward solution. We do this for the following reasons

i. Studies of dipole localization accuracy using BEM (Akalin Acar and Makeig 2013; Fuchs, Wagner, and Kastner 2007), are relatively less common than those using FEM. The use of BEM-FMM allows us to test much higher mesh resolutions than the ones using conventional BEM.
ii. BEM models can handle a much larger mesh resolution than FEM. This is because FEM requires volumetric meshes, whereas BEM only requires a triangular mesh for the tissue boundaries. On the other hand, the major disadvantage of BEM against FEM is that, although it is possible to incorporate anisotropy to BEM (Olivi, Papadopoulo, and Clerc 2011), most implementations of BEM lack this feature.
iii. Prescribing arbitrary point dipoles — which are a common mathematical model for the simultaneous firing of a large number of neurons (Hämäläinen et al. 1993) — is an easy task in BEM. On the other hand, modeling point dipoles in FEM is a more challenging task: several approaches are available (Schimpf, Ramon, and Haueisen 2002), but using them would involve an additional error estimation— see also the St. Venant approach used in (Buchner et al. 1997; Vorwerk, Aydin, et al. 2019).

Our paper’s organization is as follows: Section II describes the materials and methods used in our numerical experiments, Section III summarizes our results, and Section IV includes a brief discussion and interpretation. In Supplement A, we include additional tables of results.

## II. Materials and Methods

### Outline of methodology

We set up the following numerical experiment, which is carried out for a total of 15 subjects, 3 conductivity sets, 2 mesh segmentations, and 4 dipole locations:

i. We place a dipole at a location **p**_0_ approximately halfway between the `CSF-GM` and `GM-WM` tissue interfaces of our subject, and with an orientation **q**_0_ normal to the `CSF-GM` interface.
ii. We simulate a single time sample of noiseless EEG data using the *charge-based formulation of the boundary element method with fast multipole method acceleration* (BEM-FMM) and *adaptive mesh refinement* (AMR), over a high-resolution 5-layer (7-compartment) head volume conduction model.
iii. We use the simulated EEG data to perform source reconstruction with a low-resolution 3-layer head model, similar to the ones of widespread use in EEG source reconstruction. Doing so, we find the best fit for location **p**_1_, and orientation **q**_1_ provided by the FieldTrip Toolbox’s (Oostenveld et al. 2011) source localization procedures.
iv. We compute the distance ∥**p**_0_ − **p**_**1**_∥_2_ between both locations, and the angle between **q**_0_ and **q**_1_, to measure the error of the fit.

### 7-compartment models

We used the SimNIBS package (Saturnino et al. 2019) to obtain two segmentations from the T1- and T2-weighted MRI data from 15 subjects of the *Connectome Young Adult dataset* (Van Essen et al. 2012). One segmentation was obtained using the option `mri2mesh`, which performs segmentation based on *FreeSurfer* (Fischl 2012), and the other using `headreco`, which is based on SPM/CAT (Nielsen et al. 2018). The FreeSurfer and Headreco segmentations generate five main layers: *skin, skull, CSF, grey matter*, and *white matter*.

Each skin, skull, and CSF mesh from the FreeSurfer segmentations contains approximately 120 000 triangles, whereas the grey and white matter meshes contain circa 250 000 triangles. Headreco segmentations comprise approximately 150 000 triangles for skin and skull, 100 000 triangles for CSF, 300 000 triangles for grey matter, and 350 000 triangles for white matter.

Both segmentations include two additional tissue meshes, in the case of FreeSurfer, these are the *cerebellum* and *ventricles*; for Headreco these are the *eyes* and *ventricles*. One may argue that this should limit comparability between segmentations to some degree, however, the location of the two additional compartments is distant from most electrode positions. More details on the MRI data acquisition, spatial coregistration, segmentation, and mesh generation for this dataset can be found on (Htet et al. 2019) and (Van Essen et al. 2012).

We tested three tissue conductivity sets for each subject, which we labeled `IT’IS7` (Hasgall et al. 2022; Gabriel 1996), `VWB7` (Vorwerk, Wolters, and Baumgarten 2024; Vorwerk, Aydin, et al. 2019), and `SimNIBS7` (Saturnino et al. 2019). Its values can be found in Table 1, for an additional reference on the conductivity of living tissues see (Knösche and Haueisen 2022).

**Table 1:**
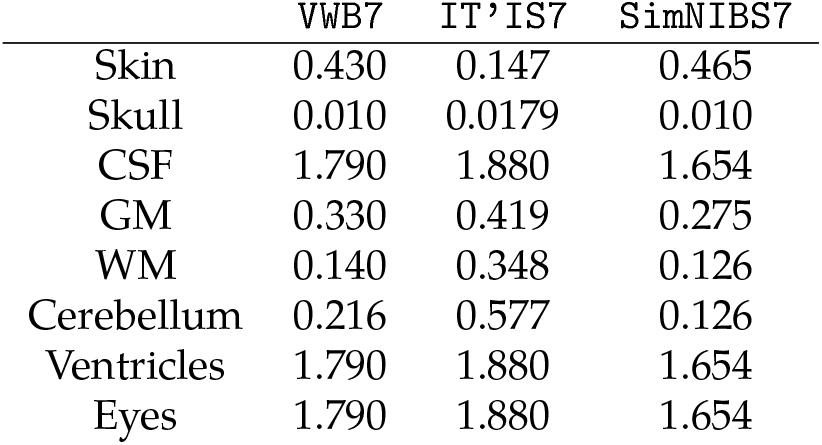
Conductivity values (S m^−1^) for the 7-compartment models.

Each of the chosen conductivity sets is optimized for different scenarios: The IT’IS7 set provides conductivity values for frequencies up to 1 MHz, `SimNIBS7` specifies conductivities adequate for TMS and TES, see also (Opitz et al. 2015), and (Wagner et al. 2004), and VWB7 has been used in several studies on the uncertainty of EEG source localization.

The only set that provides conductivity values for all the layers is `IT’IS7` (Hasgall et al. 2022), for `VWB7` there are no standard cerebellum conductivities given. We chose a 40-60 % weighted average between the conductivities of grey matter and white matter as a conductivity value for the cerebellum. `IT’IS7` assigns the same conductivity value as CSF for the eye’s vitreous humor. Hence, we assign the corresponding CSF conductivity for the eye layer in each Headreco conductivity set. We must remark, however, that the conductivity values of these additional layers are mostly irrelevant in the EEG problem since these are located far away from the scalp electrodes.

### 3-layer models

From each FreeSurfer 7-compartment head model, we created decimated (downsampled) 3-layer models using the software MeshLab (Cignoni et al. 2008). Namely, we create three new meshes labeled `SKIN, SKULL`, and `BRAIN` from the high-resolution *skin, skull*, and *CSF* FreeSurfer layers of each subject.

To obtain these new meshes, we first apply a screened Poisson surface reconstruction filter (Kazhdan and Hoppe 2013), followed by a quadratic edge collapse decimation filter with 14 000 triangles as a target. After this is done, we apply a Taubin smoothing filter (Taubin 1995).

We used these 3-layer meshes to carry out dipole source localization using the FieldTrip toolbox (Oostenveld et al. 2011). In other words, we use the 3-layer model as our inverse volume conduction model. For every choice of conductivity for the forward 5-layer model we have the corresponding 3-layer conductivities labeled `IT’IS3, VWB3`, and `SimNIBS3`, see Table 2.

**Table 2:**
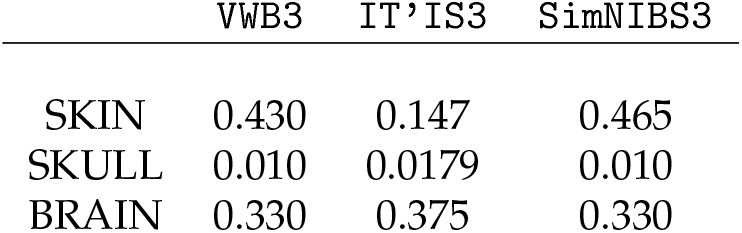
Conductivity values (S m^−1^) for the 3-layer models.

|  | VWB3 | IT'IS3 | SimNIBS3 |
| --- | --- | --- | --- |
| SKIN | 0.430 | 0.147 | 0.465 |
| SKULL | 0.010 | 0.0179 | 0.010 |
| BRAIN | 0.330 | 0.375 | 0.330 |

### Source dipole placement

We tested a total of four dipole positions for each subject. The dipole centers were chosen roughly halfway between the grey and white matter layers, and their orientation has been taken to be perpendicular to the closest grey matter region to their centers. The following are the locations chosen:

i. `dip1` — Posterior wall of the central sulcus, somatosensory cortex, tangential dipole, Figure 1.
ii. `dip2` — M1_HAND_ region, primary motor cortex, radial dipole, Figure 2
iii. `dip3` — Temporal lobe, along the Heschl’s gyri or transverse temporal gyri, Figure 3
iv. `dip4` — Medio-temporal region, Figure 4

**Figure 1:**
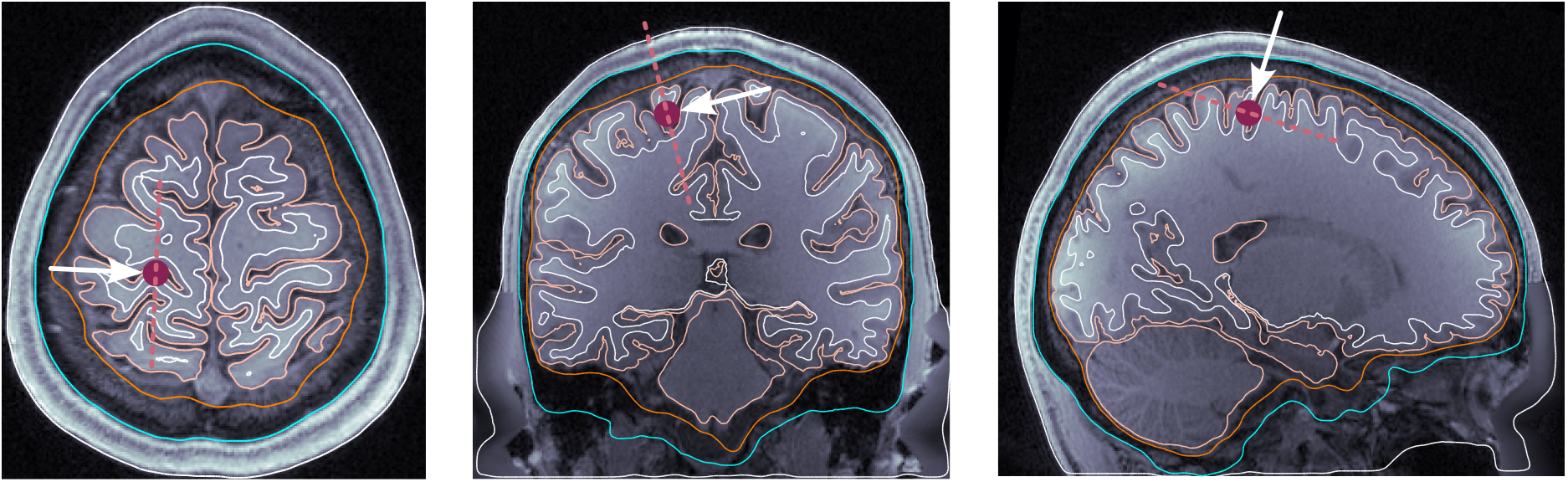
Dipole placement in the transverse, coronal, and sagittal planes, respectively, for `dip1` (posterior wall of the central sulcus) in subject 110411.

**Figure 2:**
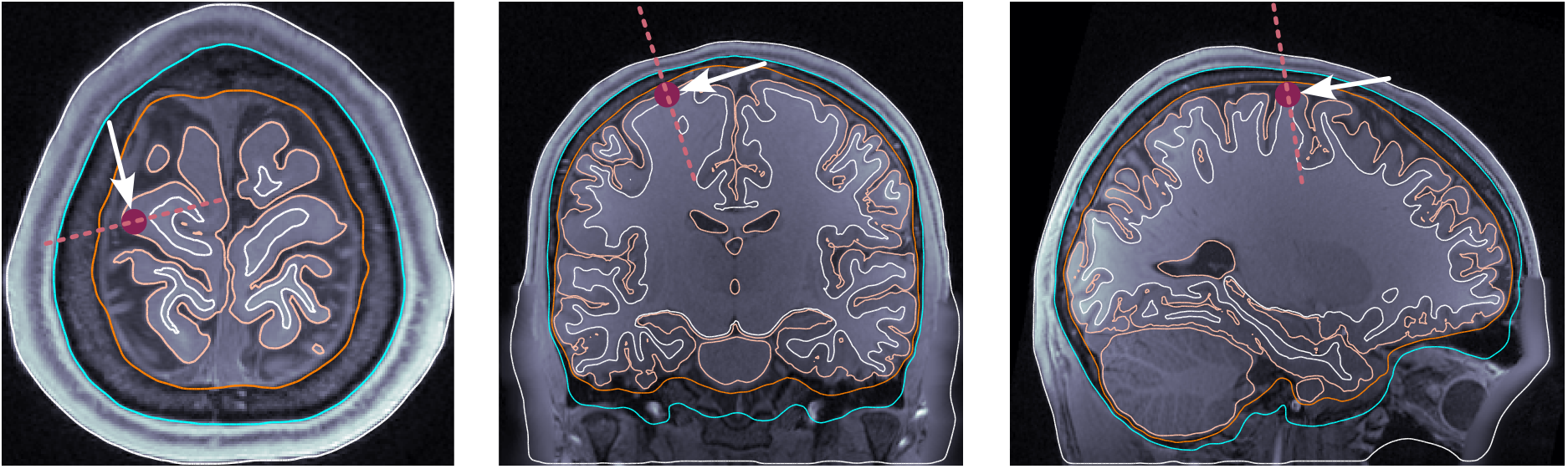
Dipole placement in the transverse, coronal, and sagittal planes, respectively, for `dip2` (M1_HAND_) in subject 130013.

**Figure 3:**
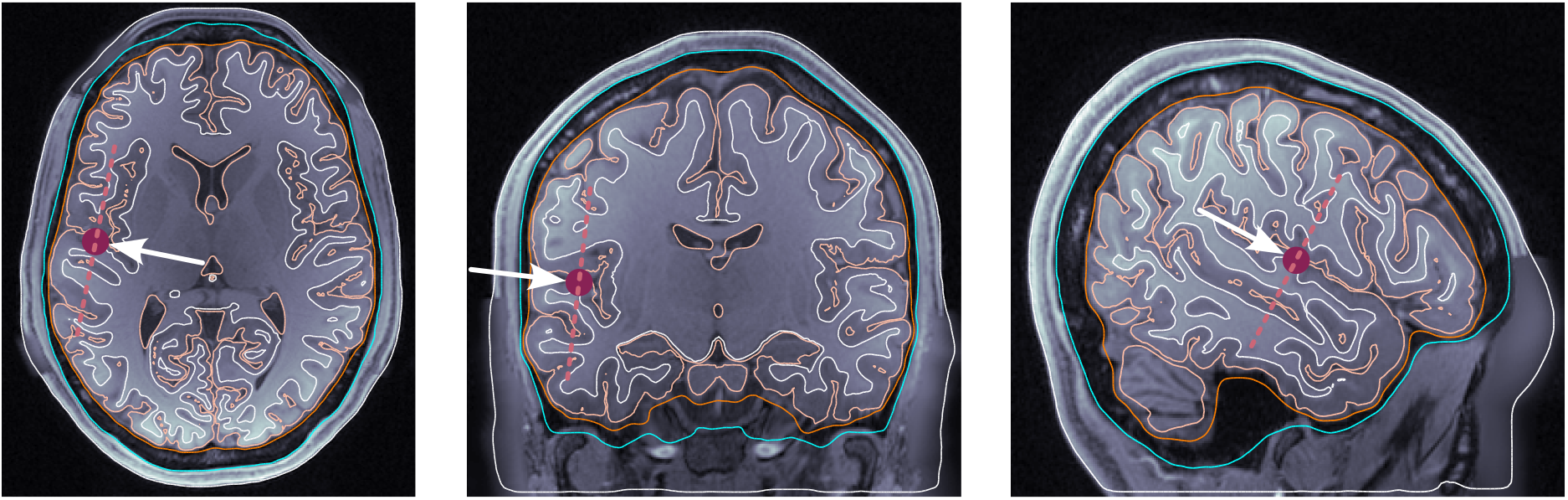
Dipole placement in the transverse, coronal, and sagittal planes, respectively, for `dip3` (temporal lobe, Heschl’s gyrus) in subject 117122.

**Figure 4:**
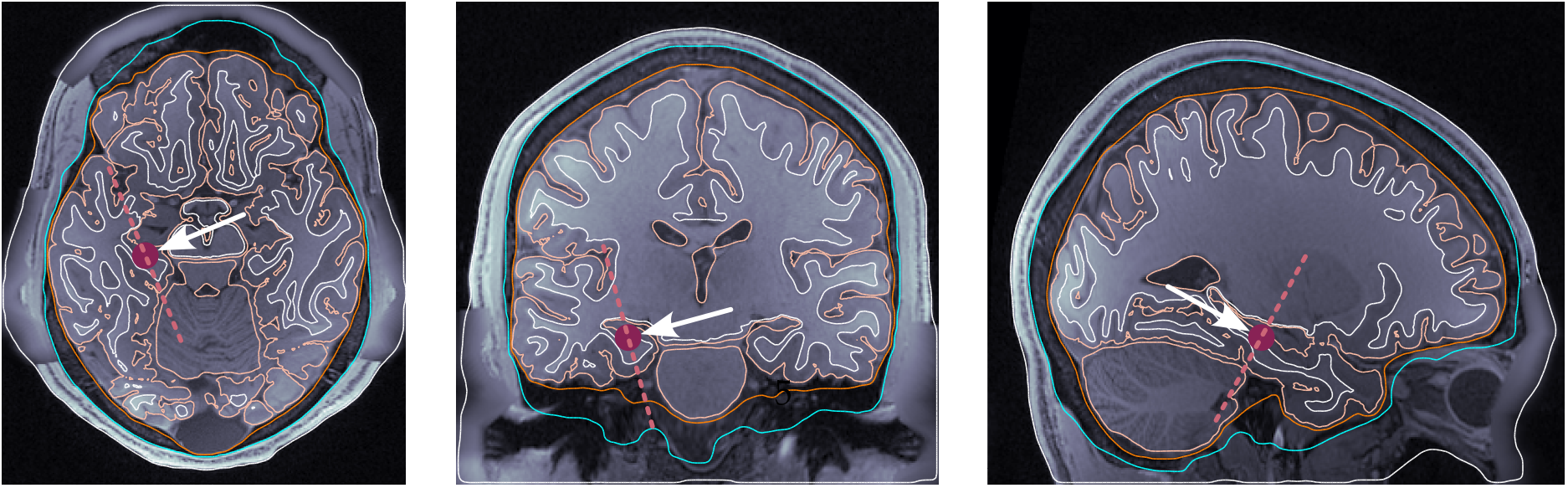
Dipole placement in the transverse, coronal, and sagittal planes, respectively, for `dip4` (medio-temporal lobe) in subject 122620.

The position `dip3` is at the superior part of the temporal cortex and has been chosen to study both auditory and language areas located at the posterior or anterior part around the same depth. The position `dip4` is relevant in epileptogenic analyses (Pittau et al. 2014).

### Electrode Placement

We placed 256 electrodes in each subject’s FreeSurfer, Headreco, and decimated FreeSurfer skin meshes, see Figure 5. The high-density 256 dry electrode system that we used is described in (Fiedler et al. 2022), see also (Graichen et al. 2015). Since our dataset (Htet et al. 2019) does not include data on the three anatomical landmarks of *nasion, left preauricular point* (LPA), or *right preauricular point* (RPA), we did a semi-manual electrode placement on each subject’s skin layer using the FieldTrip function `ft_electrode_placement`, first using the `interactive` option, followed by the `projection` option. In this way, each electrode is assigned to a triangle in the *skin* layer. We can then retrieve the electrode voltages from the BEM-FMM calculation of the skin potential. This is done by taking the value of the potential at the mesh triangle of each electrode.

**Figure 5:**
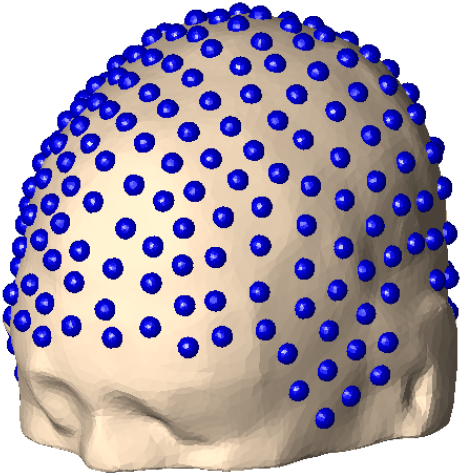
256-electrode placement on skin layer of Connectome subject 110411.

### Forward solution with BEM-FMM

We used the charge-based formulation of the BEM (Gelernter and Swihart 1964; Barnard, Duck, and Lynn 1967), together with FMM acceleration (Greengard and Rokhlin 1987).

In this formulation of the BEM forward problem, the total electric field consists of two components, an *impressed* electric field **E**^*i*^, and a *secondary* electric field **E**^*s*^. The secondary electric field arises from the *surface charge density* induced by the primary field on each surface of discontinuity of the conductivity. The boundary condition is that the normal component of the total electric current is constant across the surfaces of discontinuity. Applying the boundary condition to the total electric field **E** = **E**^*i*^ + **E**^*s*^, one derives the following integral equation for the surface charge density *ρ* over the surfaces of discontinuity *S*

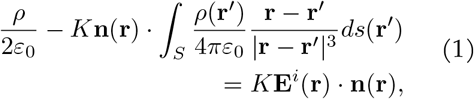

where *ε*_0_ is the permittivity of free space, **n**(**r**) is the outward normal vector to *S* at **r**, and *K* = (*σ*^−^ − *σ*^+^)*/*(*σ*^−^ + *σ*^+^) is the conductivity contrast at **r** ∈ *S*. For more details on the derivation of this equation, see (Ponasso 2023). For a model with a total of *M* facets, the discrete version of Equation (1) at the *m*-th facet is

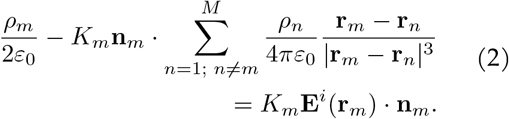

The summation term is an *n*-body computation that we accelerated using the FMM (Rokhlin 1985; Greengard and Rokhlin 1987; Beatson and Greengard 1997). In our computations, we used the FMM3D library (Askham et al. 2024).

### Adaptive mesh refinement

Adaptive mesh refinement (AMR) is a general technique that improves the convergence of the discretized solution to the FMM problem to the true analytic solution (Feischl et al. 2015). The method of AMR in the context of BEM-FMM was introduced in (Weise et al. 2022), see also (Wartman et al. 2024).

AMR consists of subdividing mesh triangles according to a cost function until a stopping criterion, based on the convergence of the solution over a region of interest, is reached. In our case, the cost function for the *m*-th triangular facet is *C*_*m*_ = |*ρ*_*m*_| · *A*_*m*_, where *ρ*_*m*_ is the charge density in the facet, and *A*_*m*_ is its area. In other words, our cost function is the total charge magnitude in each triangle. The triangles with the top 1% cost over all triangles are subdivided into four congruent triangles. After this, the surface charges are recomputed, and a new AMR step begins. The refinement process is terminated when the change in electrode voltages is less than 1% relative to the previous AMR step: Let *V*_*n*_ ∈ ℝ^256^ be the measured electrode potentials at the *n*-th AMR step, then we stop the iterative procedure at the *n*-th step provided that

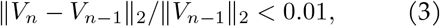

where ∥ · ∥_2_ is the 2-norm in R^256^. When we perform AMR, all layers of the model are refined except for the *skin* layer, which is our region of interest and it is used to compare the relative change from one AMR step to the next. This method is the one utilized and described in (Wartman et al. 2024). In this paper, the reader can find additional details on the AMR method we used and its integration with BEM-FMM. It also validates the method’s accuracy for the EEG dipole forward problem.

### EEG source localization

For EEG source localization, we used the FieldTrip MATLAB Toolbox (Oostenveld et al. 2011). We used the functions `ft_prepare_headmodel` with the 3-layer decimated meshes and the conductivity values in Table 2. A single-dipole source localization (with free dipole position and orientation) was carried out using non-linear optimization (Scherg 1990) via the FieldTrip function `ft_dipolefitting`. The forward solver used in the dipole fitting procedure was `bemcp` (Phillips 2001), which implements the classical surface potential formulation of the EEG forward problem (Geselowitz 1967; Sarvas 1987). We note that this forward solution does not include AMR. The reported best fit for the dipole is the least-squares approximation of the “true” EEG data by EEG data generated with the fitted dipole. The accuracy of the dipole fit is thus measured in terms of *residual variance* (RV) which is the variance of EEG data unexplained by the dipole fit. Therefore, smaller values of residual variance indicate a better fit.

We carry out two dipole fit strategies, and in every case, report the predicted dipole that has the smallest residual variance. In the first method, we start a non-linear fit with the true source dipole position as an initial value for the inverse problem (Scherg 1990). In the second method, we prepare a grid with a resolution of 5 mm using the function `ft_prepare_sourcemodel`, and a grid search is performed from every grid position within the skull. The best initial position in the grid is then used as the initial point of a non-linear fit.

## III. Results

### Summary of results

When AMR is used, the average distance error in source localization (across all models, subjects, and dipole positions we tested) caused by the discrepancy in charge-based BEM-FMM and potential-based BEM is ∼1 mm. To this error, an average of ∼3 mm distance error is added by incorporating intracortical compartments. Finally, if AMR is not used, an average of ∼4 mm distance error is added due to numerical inaccuracies. Below, we elaborate on these results.

### 3-layer model comparison

FieldTrip does not use the charge-based BEM-FMM in its forward engine, instead, it uses the classical potential-based formulation of the BEM (Geselowitz 1967; Sarvas 1987). Because of this, we made a model comparison to estimate the baseline accuracy of FieldTrip’s potential-based 3-layer inverse model (without AMR) under the assumption that the forward charge-based BEM-FMM (with and without AMR) solution calculated using the 3-layer model is ground truth. This is a necessary step since there is no analytic solution available, and knowing the discrepancy due to model differences we will be able to better assess the decrease in accuracy of dipole fitting due to the inclusion of additional layers.

First, we calculated the potentials in the entire `SKIN` layer using a forward 3-layer model with charge-based BEM-FMM and AMR. Then, we retrieved readings for the electrode potentials from this calculation using the potential value at the `SKIN` triangle corresponding to each electrode position. With these electrode voltages, we generated a raw EEG data set (compatible with FieldTrip software) consisting of a single time sample.

Finally, we performed EEG source reconstruction from the simulated EEG using a FieldTrip 3-layer volume conduction model (which does not use AMR in the forward solution) with the same conductivity values used in the forward model. See Figure 6 for a diagrammatic representation of the 3-layer model comparison process.

**Figure 6:**
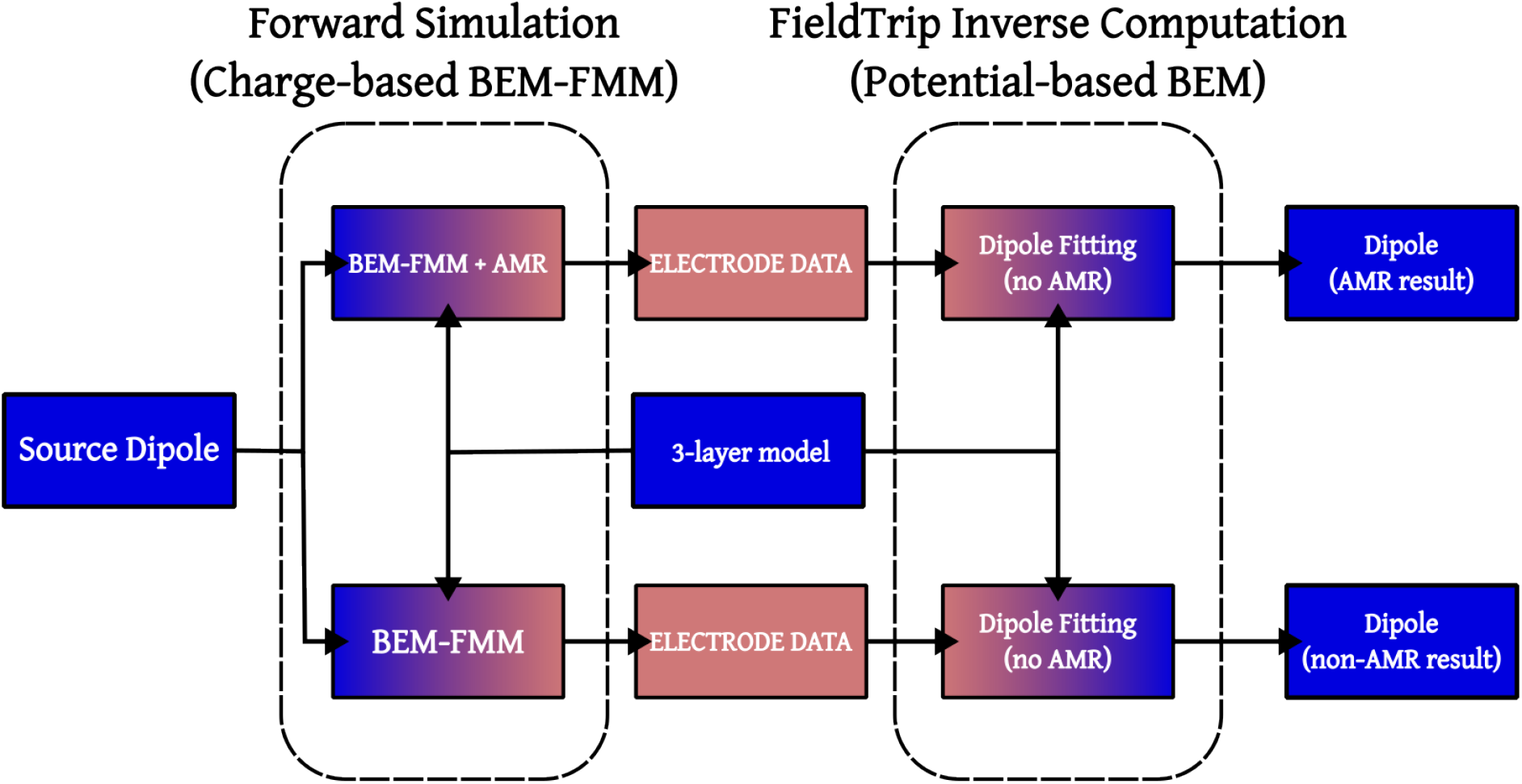
A flowchart for the 3-layer model comparison process. The blue shade represents geometric information, the pink shade represents electromagnetic information. Given a single source dipole and a 3-layer model (consisting of meshes and a tissue conductivity set), we produce two dipole fits: one corresponding to an AMR forward solution, and one corresponding to a non-AMR one.

When AMR is used, the FieldTrip inverse model retrieves the source dipole position with an average distance error of ∼1 mm (over all dipoles, subjects, and models), and an average angle error of 0.75°. If AMR is not used in the forward solution, then the average distance error of the fit is approximately 2.5 mm, with an orientation error of ∼ 2°. Detailed tables with 3-layer model comparison values can be found in Supplement A.

### 5-layer vs 3-layer results

We generated a forward BEM-FMM solution from each dipole location and each 5-layer model with AMR and without AMR and assumed that these solutions are ground truth. Then, we performed two dipole fits (one for the AMR solution and one for the non-AMR solution) using the 3-layer model of the corresponding conductivity as an inverse model (e.g. when using `IT’IS7` as the conductivity set for our forward computations, we use `IT’IS3` as the conductivity set for the inverse computation), see Figure 7. We measured the distance error and orientation error in each case.

**Figure 7:**
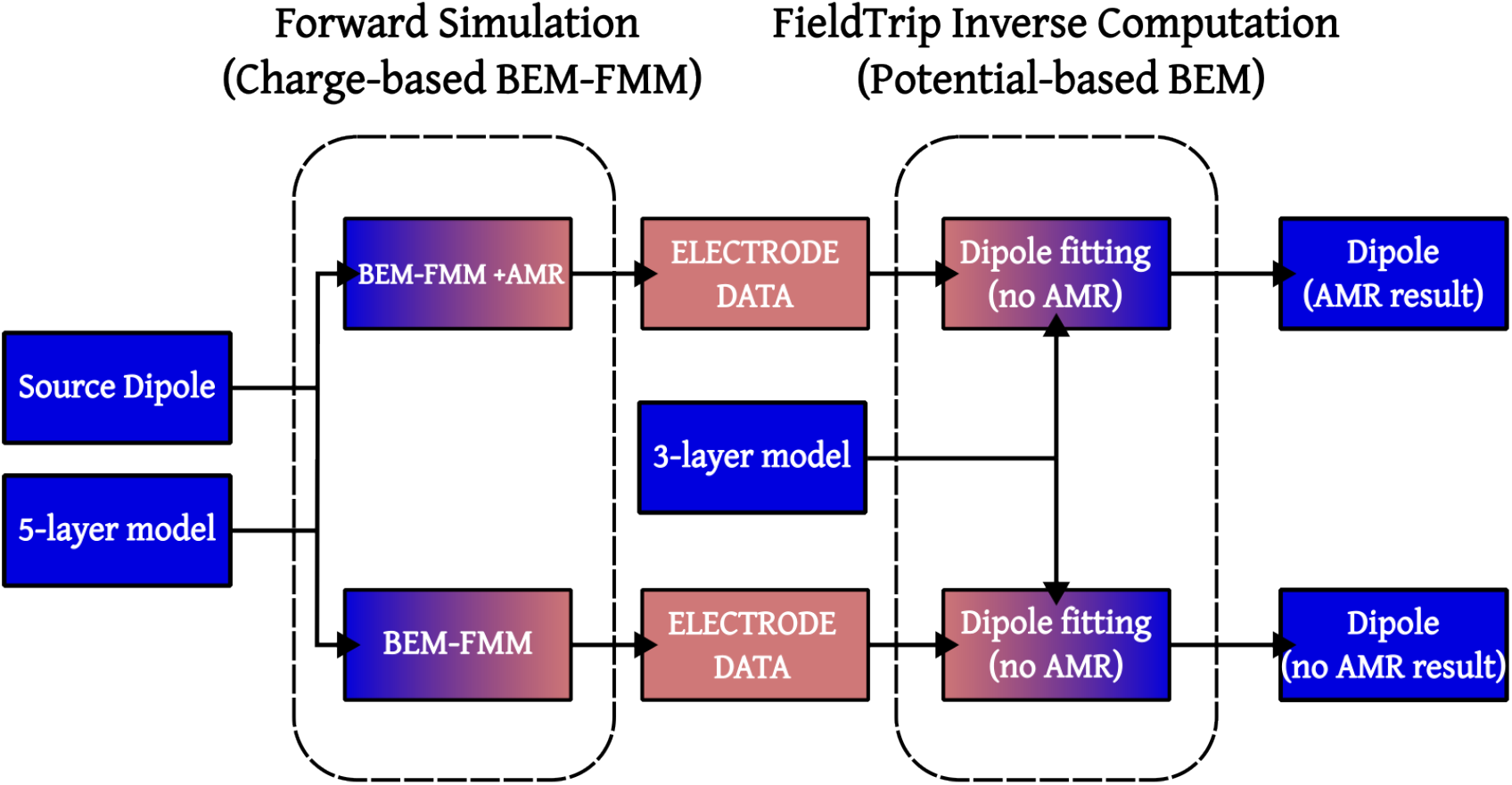
A flowchart for the performance study of 3-layer inverse models using 5-layer forward models. Given a single source dipole and a 5-layer model, we produce two dipole fits using an inverse 3-layer model: one corresponding to an AMR forward solution, and one corresponding to a non-AMR one.

When AMR is used, we obtain an average distance error among (across all dipole positions and subjects) of ∼4 mm (∼1 mm std.) using FreeSurfer meshes and an average distance error of ∼4 mm (∼2.5 mm std.) using Headreco meshes. The error in dipole orientation is more pronounced, with ∼15° average (∼10° std.) for FreeSurfer models, and ∼20° average (∼20° std.) for Headreco models. See Table 3 for dipole-wise AMR results.

**Table 3:**
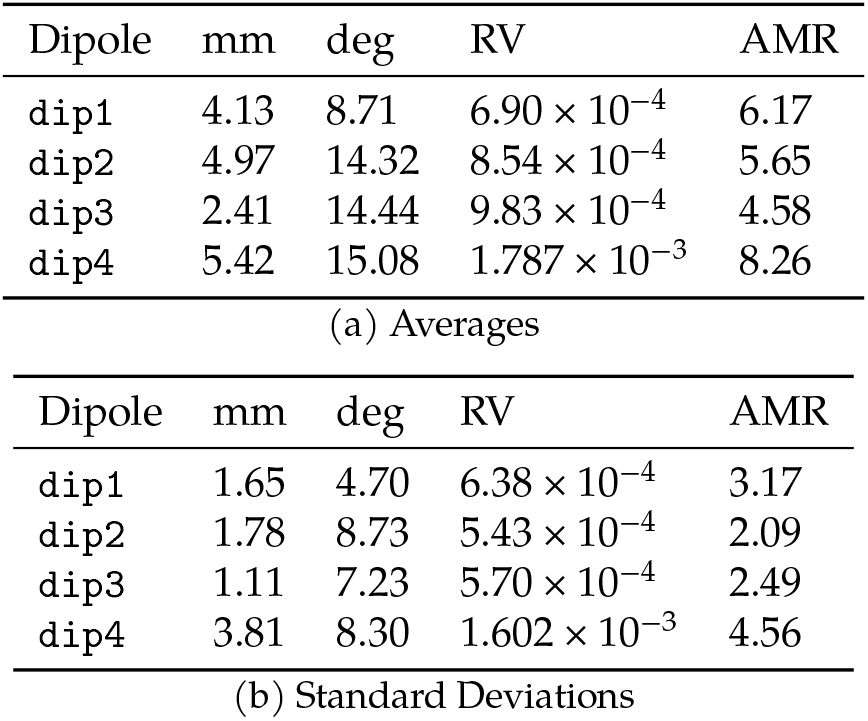
AMR results per dipole position for all 5-layer models. The column AMR denotes the total number of AMR steps in the forward solution.

| Dipole | mm | deg | RV | AMR |
| --- | --- | --- | --- | --- |
| dip1 | 4.13 | 8.71 | $6.90 \times 10^{-4}$ | 6.17 |
| dip2 | 4.97 | 14.32 | $8.54 \times 10^{-4}$ | 5.65 |
| dip3 | 2.41 | 14.44 | $9.83 \times 10^{-4}$ | 4.58 |
| dip4 | 5.42 | 15.08 | $1.787 \times 10^{-3}$ | 8.26 |
| (a) Averages |  |  |  |  |
| Dipole | mm | deg | RV | AMR |
| dip1 | 1.65 | 4.70 | $6.38 \times 10^{-4}$ | 3.17 |
| dip2 | 1.78 | 8.73 | $5.43 \times 10^{-4}$ | 2.09 |
| dip3 | 1.11 | 7.23 | $5.70 \times 10^{-4}$ | 2.49 |
| dip4 | 3.81 | 8.30 | $1.602 \times 10^{-3}$ | 4.56 |
| (b) Standard Deviations |  |  |  |  |

On the other hand, if AMR is not used, the forward solution has a large numerical error which causes worse dipole fitting results. The average distance error per dipole is ∼8 mm (∼5.5 mm std.), and the average orientation error is ∼28° (∼28° std.), see Table 4

For additional tables containing data related to the 3-layer model comparison, and model-wise results for 3-layer versus 5-layer models with and without AMR, see Supplement A.

**Table 4:** Non-AMR results per dipole for all 5-layer models.

| Dipole | mm | deg | RV |
| --- | --- | --- | --- |
| dip1 | 9.47 | 24.84 | $6.129 \times 10^{-3}$ |
| dip2 | 6.88 | 19.63 | $2.544 \times 10^{-3}$ |
| dip3 | 3.97 | 18.69 | $2.668 \times 10^{-3}$ |
| dip4 | 10.04 | 50.04 | $6.500 \times 10^{-3}$ |
| (a) Averages |  |  |  |
| Dipole | mm | deg | RV |
| dip1 | 8.50 | 17.92 | $5.543 \times 10^{-3}$ |
| dip2 | 4.60 | 30.58 | $2.612 \times 10^{-3}$ |
| dip3 | 2.94 | 11.99 | $3.004 \times 10^{-3}$ |
| dip4 | 5.84 | 50.73 | $5.788 \times 10^{-3}$ |
| (b) Standard deviations |  |  |  |

## IV. Discussion

### Summary of main findings

Dipole fitting with a low-resolution 3-layer model gives excellent results with noiseless EEG data simulated using a BEM-FMM 5-layer model with AMR. The additional accuracy in dipole fitting that may be gained by using a forward 5-layer model in the dipole fitting algorithm would be lost if the numerical error in the BEM solution calculation cannot be controlled. This requires additional techniques, such as the AMR method described in this paper.

### Dipole fittings

The average worst dipole location reconstruction result is that of dipole `dip4`, which is perhaps to be expected since its location in the medio-temporal region (Figure 4) is the deepest of all those we considered. More surprisingly, the position of `dip2` (a radial dipole located in the M1_HAND_ region) was the one that gave the second-to-worst results. The reason may be that the 3-layer models do not include CSF, and the closer proximity of the dipole to the highly conductive CSF layer creates a larger discrepancy between the 5- and 3-layer models. However, one should not over-interpret these results, as more tests for radial dipoles with varying thicknesses of neighboring CSF layers may be needed before a conclusion can be made.

### Advantages and shortcomings of the 3-layer model

The residual variances we obtain (circa 0.0015 on average) are extremely small, which indicates that the nonlinear dipole fitting using the 3-layer model explains most of the EEG data variance with the predicted positions. This indicates that these are good-quality fits. In particular, when we use AMR, the discrepancies to the original dipole positions and orientations are only caused by the difference in model complexity (5-layers vs 3-layers). If AMR is not used, then the errors are due to the combined effect of model-complexity difference and the numerical error caused by the short distance between layers, or the proximity of the dipole singularity to the triangle meshes.

Given this, we find that the 3-layer model retrieves our chosen dipole positions with a very reasonable accuracy of approximately 1-5 mm. However, the reconstruction of the dipole orientation is not as precise. Since the particular dipole location arises as a “center of mass” from the region of activity, the orientation may be more useful for determining the precise location of activity in the cortex in practical scenarios, see (Ebersole and Hawes-Ebersole 2007).

We conclude that the 3-layer model remains a very powerful tool for source localization, but we advise that end users of dipole localization software exercise caution when interpreting the results. The effects of noise and the possibility of multiple dipoles being the source of the signal were not explored in the present study.

### Advantages and challenges of the 5-layer model

Inclusion of the grey and white matter tissues is a major factor in the variation of dipole orientation, see Figure 8, this is in agreement with the findings in (Vorwerk, Aydin, et al. 2019).

**Figure 8:**
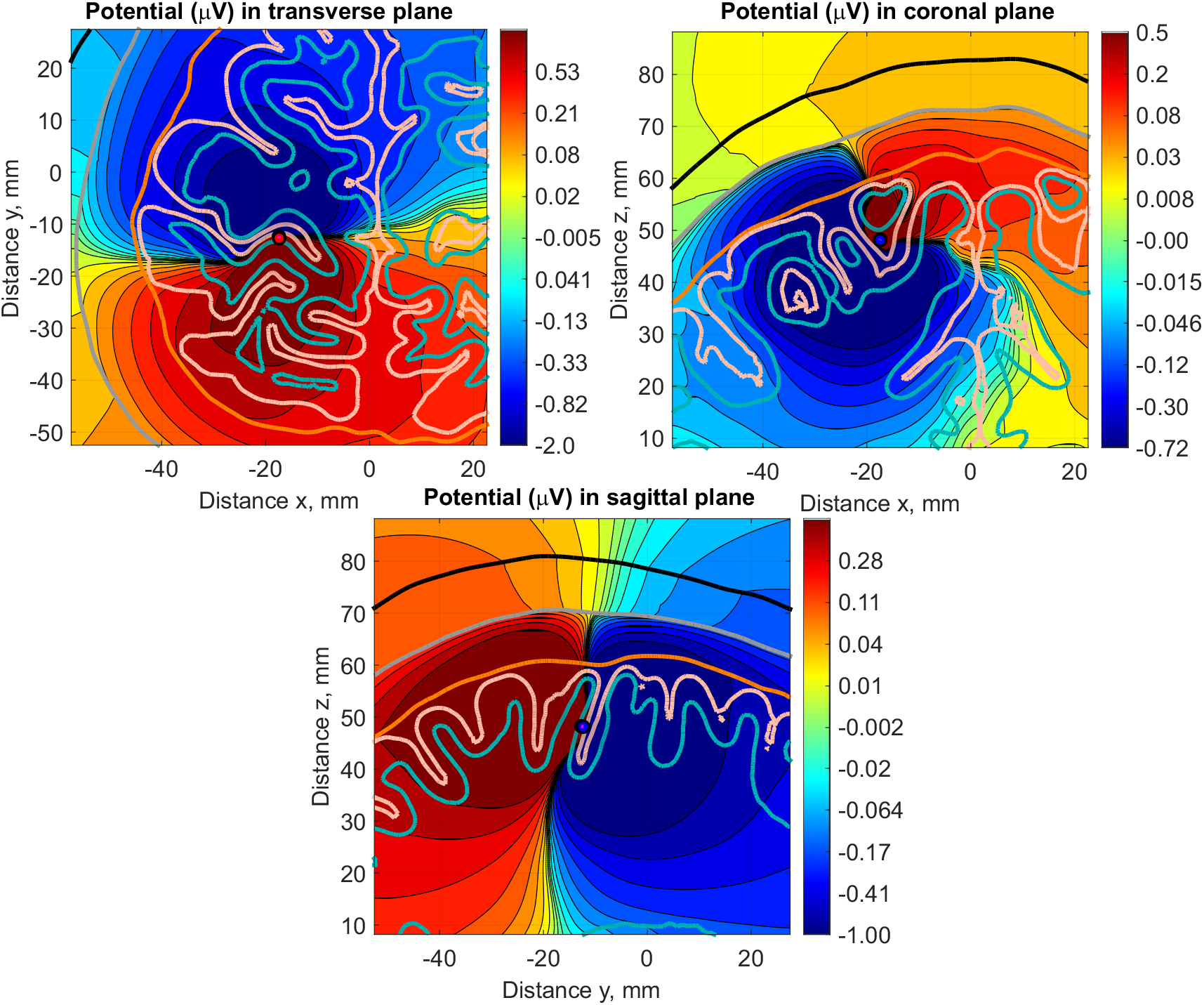
Volumetric plots in logarithmic scale of the electric potential (μV) around the source dipole `dip1` in subject 110411 —transverse, coronal, and sagittal views. Observe the symmetry break with respect to the axes of the dipole already happening in the intracortical compartments.

Model-wise, the conductivity set that produces the best agreement between the 3-layer and 5-layer models is `IT’IS`. A possible explanation is that this set has the lowest conductivity contrast between grey matter and CSF. We also observe (see Supplement A) that the dipole fits for the forward data generated with Headreco models have higher variability than those fits using the data from FreeSurfer models (both in terms of location and orientation). The use of AMR should eliminate the discrepancies caused by the different number of triangles in the initial layers, so a possible explanation for the different variabilities is that Headreco meshes use a more realistic skull layer than the FreeSurfer meshes. However, a formal assessment of the effect of mesh uncertainty would require a more detailed analysis using techniques similar to those in (Vorwerk, Aydin, et al. 2019; Vorwerk, Wolters, and Baumgarten 2024).

The results of Table 3 contrasted to Table 4 indicate that it is necessary to incorporate some form of AMR to have accurate forward computations and, in turn, an accurate dipole fitting based on 5-layer BEM models. Certain dipole positions require some layers to be refined up to 12 times before we can reduce the electrode error sufficiently. For this reason, a practical implementation of dipole fitting using BEM-FMM would require an improvement in AMR speed. This is in contrast to 3-layer models: the lack of intracortical compartments implies that the dipolar sources will be typically well-separated from the model meshes, and thus the error caused by the lack of AMR is much lower.

Modeling EEG noise can be achieved by placing dipoles at random positions, and with random strengths, all over the space between the `CSF-GM` and `GM-WM` interfaces. This task can be easily achieved with FMM-LU (Sushnikova et al. 2023) in the 3-layer model. However, FMM-LU is incompatible with AMR, and severe numerical inaccuracies can arise in the 5-layer model due to the proximity of the dipolar sources to the mesh triangles of the white and grey matter layers. Previous studies have suggested that the addition of noise mainly influences the source localization accuracy of deep sources (Whittingstall et al. 2003)

## Acknowledgements

The authors would like to thank Prof. Dr. Carsten Wolters for his help regarding conductivity values for the 3-layer model. GNP, RMcS, WAW, PL, GMN, and SNM were supported by the NIBIB grant R01EB035484, and NIMH grant R01MH130490. TR was partially supported by the NINDS grant 1R01NS126337. KW was partially supported by the BMBF grant: 01GQ2201. JH received funding from the German Federal Ministry of Education and Research (BMBF) grant Dry-Pole (01GQ2304A) and the Free State of Thuringia (2018 IZN 004), co-financed by the European Union under the European Regional Development Fund (ERDF).

## Notes

### Competing Interest Statement

The authors have declared no competing interest.

