## Supplement A for "Accuracy of dipole source reconstruction in the 3-layer BEM model against the 5-layer BEM-FMM model"

### Supplement A: Tables

#### 1 Tables of results per dipole position

Here we include tables with results per dipole separated among those experiments using forward simulated data from FreeSurfer and Headreco segmentation meshes. We also include the 3-layer model comparison data, where the forward solution has been computed using the 3-shell decimated models. Two important observations can be made:

(i) The variability of dipole fits from EEG data generated with Headreco models is larger than that generated with FreeSurfer models.

(ii) The EEG data generated without AMR has numerical errors, reflected on the fitting results with larger errors in the dipole fit, and higher variabilities.

##### 1.1 FreeSurfer segmentation results

Table 1: AMR — FreeSurfer.
Forward model: FreeSurfer with 7 high-resolution compartments computed with BEM-FMM+AMR. Inverse model: Decimated FreeSurfer shells with 3 compartments of circa 14 000 triangles.

| Dipole | Dist-mm | Angle-deg | Residual-Variance | Total-AMR-steps |
|---|---|---|---|---|
| dip1 | 4.37 | 8.70 | $5.96 \times 10^{-4}$ | 5.56 |
| dip2 | 4.38 | 14.20 | $5.49 \times 10^{-4}$ | 5.64 |
| dip3 | 2.47 | 14.52 | $9.59 \times 10^{-4}$ | 4.29 |
| dip4 | 4.12 | 18.07 | $2.474 \times 10^{-3}$ | 8.60 |

(a) Means

| Dipole | Dist-mm | Angle-deg | Residual-Variance | Total-AMR-steps |
|---|---|---|---|---|
| dip1 | 1.55 | 4.50 | $8.22 \times 10^{-4}$ | 2.49 |
| dip2 | 1.27 | 7.51 | $3.10 \times 10^{-4}$ | 2.05 |
| dip3 | 1.07 | 7.16 | $6.75 \times 10^{-4}$ | 1.85 |
| dip4 | 1.87 | 9.27 | $1.921 \times 10^{-3}$ | 4.21 |

(b) Standard Deviations

The following table includes the results of the dipole fitting without using AMR in the computation of the forward solution. Notice the dramatic decrease in fitting accuracy, and the higher variability, both of which are caused by numerical inaccuracies.

Table 2: non-AMR — FreeSurfer. Forward model: FreeSurfer with 7 high-resolution compartments computed with BEM-FMM (no AMR). Inverse model: Decimated FreeSurfer shells with 3 compartments of circa 14 000 triangles.

| Dipole | Dist-mm | Angle-deg | Residual-Variance |
|---|---|---|---|
| `dip1` | 8.45 | 22.66 | $4.918 \times 10^{-3}$ |
| `dip2` | 5.38 | 13.72 | $1.496 \times 10^{-3}$ |
| `dip3` | 4.29 | 19.80 | $2.913 \times 10^{-3}$ |
| `dip4` | 14.78 | 81.03 | $9.998 \times 10^{-3}$ |

(a) means

| Dipole | Dist-mm | Angle-deg | Residual-Variance |
|---|---|---|---|
| `dip1` | 8.27 | 15.09 | $3.955 \times 10^{-3}$ |
| `dip2` | 2.65 | 8.14 | $1.534 \times 10^{-3}$ |
| `dip3` | 3.18 | 13.59 | $2.833 \times 10^{-3}$ |
| `dip4` | 3.71 | 55.42 | $4.902 \times 10^{-3}$ |

(b) Standard Deviations

#### 1.2 Headreco segmentation results

Table 3: AMR — Headreco

| Dipole | Dist-mm | Angle-deg | Residual-Variance | Total-AMR-steps |
|---|---|---|---|---|
| `dip1` | 3.91 | 8.72 | $7.84 \times 10^{-4}$ | 6.80 |
| `dip2` | 5.57 | 14.45 | $1.160 \times 10^{-3}$ | 5.67 |
| `dip3` | 2.37 | 14.36 | $1.008 \times 10^{-3}$ | 4.89 |
| `dip4` | 6.73 | 12.10 | $1.099 \times 10^{-3}$ | 7.93 |

(a) AMR means — Headreco

| Dipole | Dist-mm | Angle-deg | Residual-Variance | Total-AMR-steps |
|---|---|---|---|---|
| `dip1` | 1.73 | 4.95 | $3.60 \times 10^{-4}$ | 3.66 |
| `dip2` | 2.03 | 9.90 | $5.56 \times 10^{-4}$ | 2.17 |
| `dip3` | 1.16 | 7.39 | $4.49 \times 10^{-4}$ | 3.00 |
| `dip4` | 4.74 | 5.93 | $7.31 \times 10^{-4}$ | 4.92 |

(b) AMR Standard Deviations — Headreco

Table 4: non-AMR — Headreco

| Dipole | Dist-mm | Angle-deg | Residual-Variance |
|---|---|---|---|
| dip1 | 10.49 | 27.01 | $7.340 \times 10^{-3}$ |
| dip2 | 8.37 | 25.54 | $3.592 \times 10^{-3}$ |
| dip3 | 3.66 | 17.58 | $2.423 \times 10^{-3}$ |
| dip4 | 5.31 | 19.05 | $3.002 \times 10^{-3}$ |

(a) non-AMR means — Headreco

| Dipole | Dist-mm | Angle-deg | Residual-Variance |
|---|---|---|---|
| dip1 | 8.71 | 20.30 | $6.596 \times 10^{-3}$ |
| dip2 | 5.58 | 41.87 | $3.033 \times 10^{-3}$ |
| dip3 | 2.67 | 10.16 | $3.179 \times 10^{-3}$ |
| dip4 | 3.09 | 13.05 | $4.325 \times 10^{-3}$ |

(b) non-AMR Standard Deviations — Headreco

#### 1.3 3-layer model comparison results

Table 5: AMR results per dipole — 3-layer model comparison

| Dipole | Dist-mm | Angle-deg | Residual-Variance | Total-AMR-steps |
|---|---|---|---|---|
| dip1 | 1.02 | 0.57 | $5.5 \times 10^{-5}$ | 2.24 |
| dip2 | 2.40 | 1.41 | $3.23 \times 10^{-4}$ | 3.31 |
| dip3 | 0.39 | 0.60 | $2.0 \times 10^{-5}$ | 2.96 |
| dip4 | 0.42 | 0.42 | $1.3 \times 10^{-5}$ | 2.78 |

(a) AMR Averages per dipole — 3-layer model comparison

| Dipole | Dist-mm | Angle-deg | Residual-Variance | Total-AMR-steps |
|---|---|---|---|---|
| dip1 | 0.27 | 0.36 | $2.8 \times 10^{-5}$ | 0.43 |
| dip2 | 0.67 | 1.32 | $2.62 \times 10^{-4}$ | 1.02 |
| dip3 | 0.16 | 0.44 | $1.2 \times 10^{-5}$ | 1.15 |
| dip4 | 0.19 | 0.32 | $9 \times 10^{-6}$ | 1.02 |

(b) AMR Standard deviations per dipole — 3-layer model comparison

Table 6: No-AMR Averages per dipole — 3-layer model comparison

| Dipole | Dist-mm | Angle-deg | Residual-Variance |
|---|---|---|---|
| dip1 | 1.59 | 2.60 | $2.93 \times 10^{-4}$ |
| dip2 | 2.08 | 2.77 | $9.10 \times 10^{-4}$ |
| dip3 | 3.06 | 1.62 | $1.79 \times 10^{-4}$ |
| dip4 | 3.04 | 1.18 | $2.70 \times 10^{-4}$ |

(a) No-AMR Averages per dipole — 3-layer model comparison

| Dipole | Dist-mm | Angle-deg | Residual-Variance |
|---|---|---|---|
| dip1 | 1.00 | 2.57 | $2.50 \times 10^{-4}$ |
| dip2 | 0.88 | 2.71 | $5.56 \times 10^{-4}$ |
| dip3 | 2.10 | 1.77 | $1.50 \times 10^{-4}$ |
| dip4 | 2.18 | 0.94 | $3.43 \times 10^{-4}$ |

(b) No-AMR Standard deviations per dipole — 3-layer model comparison

#### 1.4 Combined AMR results: FreeSurfer+Headreco

Table 7: AMR — all 7-shell models

| Dipole | Dist-mm | Angle-deg | Residual-Variance | Total-AMR-steps |
|---|---|---|---|---|
| dip1 | 4.14 | 8.71 | $6.90 \times 10^{-4}$ | 6.18 |
| dip2 | 4.98 | 14.33 | $8.54 \times 10^{-4}$ | 5.66 |
| dip3 | 2.42 | 14.44 | $9.83 \times 10^{-4}$ | 4.59 |
| dip4 | 5.42 | 15.08 | $1.787 \times 10^{-3}$ | 8.27 |

(a) AMR means — all 7-shell models

| Dipole | Dist-mm | Angle-deg | Residual-Variance | Total-AMR-steps |
|---|---|---|---|---|
| dip1 | 1.65 | 4.70 | $6.38 \times 10^{-4}$ | 3.17 |
| dip2 | 1.78 | 8.74 | $5.43 \times 10^{-4}$ | 2.10 |
| dip3 | 1.11 | 7.24 | $5.70 \times 10^{-4}$ | 2.50 |
| dip4 | 3.82 | 8.30 | $1.602 \times 10^{-3}$ | 4.56 |

(b) AMR Standard Deviations — all 7-shell models

#### 1.5 Combined Non-AMR results: FreeSurfer+Headreco

Table 8: Non-AMR results per dipole — all 7-shell models

| Dipole | Dist-mm | Angle-deg | Residual-Variance |
|---|---|---|---|
| `dip1` | 9.47 | 24.84 | $6.129 \times 10^{-3}$ |
| `dip2` | 6.88 | 19.63 | $2.544 \times 10^{-3}$ |
| `dip3` | 3.97 | 18.69 | $2.668 \times 10^{-3}$ |
| `dip4` | 10.04 | 50.04 | $6.500 \times 10^{-3}$ |

(a) Non-AMR means — all 7-shell models

| Dipole | Dist-mm | Angle-deg | Residual-Variance |
|---|---|---|---|
| `dip1` | 9.47 | 24.84 | $6.129 \times 10^{-3}$ |
| `dip2` | 6.88 | 19.63 | $2.544 \times 10^{-3}$ |
| `dip3` | 3.97 | 18.69 | $2.668 \times 10^{-3}$ |
| `dip4` | 10.04 | 50.04 | $6.500 \times 10^{-3}$ |

(b) Non-AMR standard deviations — all 7-shell models

### 2 Tables of results per model and dipole

#### 2.1 FreeSurfer segmentation results

Table 9: AMR — FreeSurfer

| Model | Dipole | Dist-mm | Angle-deg | Residual-Variance | Total-AMR-steps |
|---|---|---|---|---|---|
| SimNIBS7 | dip1 | 4.77 | 10.41 | $5.04 \times 10^{-4}$ | 6.33 |
| SimNIBS7 | dip2 | 4.66 | 15.39 | $4.92 \times 10^{-4}$ | 6.07 |
| SimNIBS7 | dip3 | 2.88 | 17.42 | $9.64 \times 10^{-4}$ | 4.93 |
| SimNIBS7 | dip4 | 5.00 | 22.12 | $3.38 \times 10^{-3}$ | 8.93 |
| VWB7 | dip1 | 5.06 | 9.94 | $8.37 \times 10^{-4}$ | 5.40 |
| VWB7 | dip2 | 4.93 | 17.24 | $5.84 \times 10^{-4}$ | 6.07 |
| VWB7 | dip3 | 2.96 | 16.87 | $1.01 \times 10^{-3}$ | 4.53 |
| VWB7 | dip4 | 4.93 | 20.25 | $2.81 \times 10^{-3}$ | 8.67 |
| IT'IS7 | dip1 | 3.26 | 5.75 | $4.47 \times 10^{-4}$ | 4.93 |
| IT'IS7 | dip2 | 3.57 | 9.96 | $5.70 \times 10^{-4}$ | 4.80 |
| IT'IS7 | dip3 | 1.56 | 9.28 | $9.04 \times 10^{-4}$ | 3.40 |
| IT'IS7 | dip4 | 2.43 | 11.82 | $1.23 \times 10^{-3}$ | 8.20 |

(a) AMR means — FreeSurfer

| Model | Dipole | Dist-mm | Angle-deg | Residual-Variance | Total-AMR-steps |
|---|---|---|---|---|---|
| SimNIBS7 | dip1 | 1.19 | 4.44 | $3.07 \times 10^{-4}$ | 2.66 |
| SimNIBS7 | dip2 | 1.05 | 7.42 | $2.53 \times 10^{-4}$ | 2.05 |
| SimNIBS7 | dip3 | 0.94 | 5.47 | $7.79 \times 10^{-4}$ | 1.87 |
| SimNIBS7 | dip4 | 1.84 | 9.99 | $1.94 \times 10^{-3}$ | 5.96 |
| VWB7 | dip1 | 1.89 | 4.21 | $1.38 \times 10^{-3}$ | 2.90 |
| VWB7 | dip2 | 1.50 | 7.66 | $2.99 \times 10^{-4}$ | 2.02 |
| VWB7 | dip3 | 1.01 | 6.48 | $8.05 \times 10^{-4}$ | 1.88 |
| VWB7 | dip4 | 1.45 | 8.73 | $2.13 \times 10^{-3}$ | 4.22 |
| IT'IS7 | dip1 | 0.75 | 3.48 | $1.69 \times 10^{-4}$ | 1.71 |
| IT'IS7 | dip2 | 0.77 | 5.73 | $3.77 \times 10^{-4}$ | 1.93 |
| IT'IS7 | dip3 | 0.64 | 6.71 | $4.11 \times 10^{-4}$ | 1.55 |
| IT'IS7 | dip4 | 0.91 | 5.44 | $7.98 \times 10^{-4}$ | 1.42 |

(b) AMR Standard Deviations — FreeSurfer

Table 10: non-AMR — FreeSurfer

| Model | Dipole | Dist-mm | Angle-deg | Residual-Variance |
|---|---|---|---|---|
| SimNIBS7 | dip1 | 9.55 | 26.10 | $4.63 \times 10^{-3}$ |
| SimNIBS7 | dip2 | 5.89 | 15.25 | $1.75 \times 10^{-3}$ |
| SimNIBS7 | dip3 | 5.52 | 25.36 | $3.22 \times 10^{-3}$ |
| SimNIBS7 | dip4 | 15.87 | 89.49 | $9.95 \times 10^{-3}$ |
| VWB7 | dip1 | 10.16 | 26.68 | $5.37 \times 10^{-3}$ |
| VWB7 | dip2 | 6.18 | 15.92 | $1.57 \times 10^{-3}$ |
| VWB7 | dip3 | 5.16 | 21.90 | $3.16 \times 10^{-3}$ |
| VWB7 | dip4 | 15.34 | 78.53 | $8.89 \times 10^{-3}$ |
| IT'IS7 | dip1 | 5.65 | 15.20 | $4.74 \times 10^{-3}$ |
| IT'IS7 | dip2 | 4.07 | 10.00 | $1.18 \times 10^{-3}$ |
| IT'IS7 | dip3 | 2.18 | 12.14 | $2.36 \times 10^{-3}$ |
| IT'IS7 | dip4 | 13.12 | 75.06 | $1.12 \times 10^{-2}$ |

(a) non-AMR means — FreeSurfer

| Model | Dipole | Dist-mm | Angle-deg | Residual-Variance |
|---|---|---|---|---|
| SimNIBS7 | dip1 | 9.40 | 15.46 | $2.87 \times 10^{-3}$ |
| SimNIBS7 | dip2 | 3.31 | 8.82 | $1.72 \times 10^{-3}$ |
| SimNIBS7 | dip3 | 3.57 | 17.43 | $2.31 \times 10^{-3}$ |
| SimNIBS7 | dip4 | 3.04 | 50.93 | $4.73 \times 10^{-3}$ |
| VWB7 | dip1 | 9.14 | 16.49 | $3.66 \times 10^{-3}$ |
| VWB7 | dip2 | 2.71 | 8.41 | $1.62 \times 10^{-3}$ |
| VWB7 | dip3 | 3.15 | 9.94 | $3.31 \times 10^{-3}$ |
| VWB7 | dip4 | 3.71 | 61.10 | $3.32 \times 10^{-3}$ |
| IT'IS7 | dip1 | 5.48 | 10.72 | $5.21 \times 10^{-3}$ |
| IT'IS7 | dip2 | 0.97 | 6.11 | $1.27 \times 10^{-3}$ |
| IT'IS7 | dip3 | 1.37 | 8.79 | $2.90 \times 10^{-3}$ |
| IT'IS7 | dip4 | 3.95 | 56.60 | $6.27 \times 10^{-3}$ |

(b) non-AMR Standard Deviations — FreeSurfer

#### 2.2 Headreco segmentation results

Table 11: AMR — Headreco

| Model | Dipole | Dist-mm | Angle-deg | Residual-Variance | Total-AMR-steps |
|---|---|---|---|---|---|
| SimNIBS7 | dip1 | 3.81 | 10.29 | $9.46 \times 10^{-4}$ | 7.93 |
| SimNIBS7 | dip2 | 6.18 | 16.16 | $1.26 \times 10^{-3}$ | 6.40 |
| SimNIBS7 | dip3 | 2.87 | 17.59 | $1.20 \times 10^{-3}$ | 5.60 |
| SimNIBS7 | dip4 | 9.14 | 13.93 | $1.30 \times 10^{-3}$ | 9.60 |
| VWB7 | dip1 | 3.86 | 10.14 | $9.13 \times 10^{-4}$ | 7.13 |
| VWB7 | dip2 | 6.12 | 16.35 | $1.27 \times 10^{-3}$ | 6.33 |
| VWB7 | dip3 | 2.85 | 17.72 | $1.18 \times 10^{-3}$ | 5.13 |
| VWB7 | dip4 | 8.67 | 13.50 | $1.25 \times 10^{-3}$ | 8.53 |
| IT'IS7 | dip1 | 4.06 | 5.75 | $4.92 \times 10^{-4}$ | 5.33 |
| IT'IS7 | dip2 | 4.41 | 10.86 | $9.54 \times 10^{-4}$ | 4.27 |
| IT'IS7 | dip3 | 1.39 | 7.77 | $6.44 \times 10^{-4}$ | 3.93 |
| IT'IS7 | dip4 | 2.38 | 8.87 | $7.41 \times 10^{-4}$ | 5.67 |

(a) AMR means — Headreco

| Model | Dipole | Dist-mm | Angle-deg | Residual-Variance | Total-AMR-steps |
|---|---|---|---|---|---|
| SimNIBS7 | dip1 | 2.04 | 5.36 | $3.64 \times 10^{-4}$ | 4.92 |
| SimNIBS7 | dip2 | 2.01 | 10.44 | $5.32 \times 10^{-4}$ | 2.03 |
| SimNIBS7 | dip3 | 1.09 | 6.52 | $4.28 \times 10^{-4}$ | 3.72 |
| SimNIBS7 | dip4 | 4.45 | 6.29 | $8.01 \times 10^{-4}$ | 6.15 |
| VWB7 | dip1 | 1.95 | 5.21 | $3.06 \times 10^{-4}$ | 3.18 |
| VWB7 | dip2 | 2.01 | 10.30 | $5.58 \times 10^{-4}$ | 1.91 |
| VWB7 | dip3 | 1.08 | 6.95 | $4.41 \times 10^{-4}$ | 3.09 |
| VWB7 | dip4 | 4.40 | 6.22 | $7.54 \times 10^{-4}$ | 4.79 |
| IT'IS7 | dip1 | 1.17 | 2.62 | $2.10 \times 10^{-4}$ | 1.99 |
| IT'IS7 | dip2 | 1.62 | 8.51 | $5.55 \times 10^{-4}$ | 1.98 |
| IT'IS7 | dip3 | 0.56 | 3.33 | $2.07 \times 10^{-4}$ | 1.83 |
| IT'IS7 | dip4 | 0.97 | 3.94 | $5.13 \times 10^{-4}$ | 2.53 |

(b) AMR Standard Deviations — Headreco

Table 12: non-AMR — Headreco

| Model | Dipole | Dist-mm | Angle-deg | Residual-Variance |
|---|---|---|---|---|
| SimNIBS7 | dip1 | 12.12 | 31.67 | $8.09 \times 10^{-3}$ |
| SimNIBS7 | dip2 | 10.34 | 27.00 | $4.28 \times 10^{-3}$ |
| SimNIBS7 | dip3 | 4.88 | 22.05 | $3.13 \times 10^{-3}$ |
| SimNIBS7 | dip4 | 6.07 | 21.47 | $3.70 \times 10^{-3}$ |
| VWB7 | dip1 | 11.63 | 30.62 | $7.72 \times 10^{-3}$ |
| VWB7 | dip2 | 9.69 | 27.16 | $4.03 \times 10^{-3}$ |
| VWB7 | dip3 | 4.44 | 21.40 | $2.82 \times 10^{-3}$ |
| VWB7 | dip4 | 5.84 | 20.43 | $3.47 \times 10^{-3}$ |
| IT'IS7 | dip1 | 7.73 | 18.75 | $6.21 \times 10^{-3}$ |
| IT'IS7 | dip2 | 5.10 | 22.48 | $2.46 \times 10^{-3}$ |
| IT'IS7 | dip3 | 1.66 | 9.27 | $1.32 \times 10^{-3}$ |
| IT'IS7 | dip4 | 4.03 | 15.25 | $1.83 \times 10^{-3}$ |

(a) non-AMR means — Headreco

| Model | Dipole | Dist-mm | Angle-deg | Residual-Variance |
|---|---|---|---|---|
| SimNIBS7 | dip1 | 9.38 | 21.05 | $6.58 \times 10^{-3}$ |
| SimNIBS7 | dip2 | 6.35 | 42.69 | $3.21 \times 10^{-3}$ |
| SimNIBS7 | dip3 | 2.93 | 10.53 | $3.78 \times 10^{-3}$ |
| SimNIBS7 | dip4 | 3.27 | 14.03 | $5.20 \times 10^{-3}$ |
| VWB7 | dip1 | 9.31 | 20.57 | $6.44 \times 10^{-3}$ |
| VWB7 | dip2 | 6.01 | 42.52 | $3.21 \times 10^{-3}$ |
| VWB7 | dip3 | 2.56 | 9.20 | $3.59 \times 10^{-3}$ |
| VWB7 | dip4 | 3.05 | 13.49 | $4.91 \times 10^{-3}$ |
| IT'IS7 | dip1 | 7.17 | 17.86 | $7.06 \times 10^{-3}$ |
| IT'IS7 | dip2 | 1.99 | 43.19 | $2.50 \times 10^{-3}$ |
| IT'IS7 | dip3 | 0.87 | 4.25 | $1.60 \times 10^{-3}$ |
| IT'IS7 | dip4 | 2.70 | 11.51 | $2.32 \times 10^{-3}$ |

(b) non-AMR Standard Deviations — Headreco

#### 2.3 Calibration results

Table 13: AMR — Calibration

| Model | Dipole | Dist-mm | Angle-deg | Residual-Variance | Total-AMR-steps |
|---|---|---|---|---|---|
| SimNIBS3 | dip1 | 1.14 | 0.73 | $6.61 \times 10^{-5}$ | 2.40 |
| SimNIBS3 | dip2 | 2.85 | 1.54 | $2.95 \times 10^{-4}$ | 3.87 |
| SimNIBS3 | dip3 | 0.38 | 0.85 | $2.24 \times 10^{-5}$ | 3.20 |
| SimNIBS3 | dip4 | 0.52 | 0.59 | $1.79 \times 10^{-5}$ | 3.00 |
| VWB3 | dip1 | 1.14 | 0.69 | $6.42 \times 10^{-5}$ | 2.33 |
| VWB3 | dip2 | 2.75 | 1.77 | $3.89 \times 10^{-4}$ | 3.80 |
| VWB3 | dip3 | 0.38 | 0.78 | $2.18 \times 10^{-5}$ | 3.13 |
| VWB3 | dip4 | 0.48 | 0.54 | $1.61 \times 10^{-5}$ | 2.93 |
| IT'IS3 | dip1 | 0.79 | 0.30 | $3.49 \times 10^{-5}$ | 2.00 |
| IT'IS3 | dip2 | 1.60 | 0.92 | $2.86 \times 10^{-4}$ | 2.27 |
| IT'IS3 | dip3 | 0.40 | 0.18 | $1.68 \times 10^{-5}$ | 2.53 |
| IT'IS3 | dip4 | 0.25 | 0.13 | $5.08 \times 10^{-6}$ | 2.40 |

(a) AMR means — Calibration

| Model | Dipole | Dist-mm | Angle-deg | Residual-Variance | Total-AMR-steps |
|---|---|---|---|---|---|
| SimNIBS3 | dip1 | 0.25 | 0.36 | $2.87 \times 10^{-5}$ | 0.51 |
| SimNIBS3 | dip2 | 0.36 | 1.36 | $1.77 \times 10^{-4}$ | 0.64 |
| SimNIBS3 | dip3 | 0.18 | 0.40 | $1.30 \times 10^{-5}$ | 1.26 |
| SimNIBS3 | dip4 | 0.19 | 0.32 | $8.24 \times 10^{-6}$ | 1.07 |
| VWB3 | dip1 | 0.24 | 0.36 | $2.76 \times 10^{-5}$ | 0.49 |
| VWB3 | dip2 | 0.40 | 1.67 | $3.74 \times 10^{-4}$ | 0.68 |
| VWB3 | dip3 | 0.19 | 0.38 | $1.31 \times 10^{-5}$ | 1.30 |
| VWB3 | dip4 | 0.18 | 0.28 | $7.66 \times 10^{-6}$ | 1.16 |
| IT'IS3 | dip1 | 0.13 | 0.16 | $1.58 \times 10^{-5}$ | 0.00 |
| IT'IS3 | dip2 | 0.30 | 0.67 | $1.96 \times 10^{-4}$ | 0.80 |
| IT'IS3 | dip3 | 0.12 | 0.13 | $1.05 \times 10^{-5}$ | 0.74 |
| IT'IS3 | dip4 | 0.06 | 0.05 | $2.46 \times 10^{-6}$ | 0.74 |

(b) AMR Standard Deviations — Calibration

Table 14: non-AMR — Calibration

| Model | Dipole | Dist-mm | Angle-deg | Residual-Variance |
|---|---|---|---|---|
| SimNIBS3 | dip1 | 2.22 | 3.92 | $4.41 \times 10^{-4}$ |
| SimNIBS3 | dip2 | 2.48 | 3.76 | $1.22 \times 10^{-3}$ |
| SimNIBS3 | dip3 | 4.67 | 2.46 | $2.78 \times 10^{-4}$ |
| SimNIBS3 | dip4 | 4.64 | 1.75 | $4.32 \times 10^{-4}$ |
| VWB3 | dip1 | 2.01 | 3.54 | $3.89 \times 10^{-4}$ |
| VWB3 | dip2 | 2.41 | 3.59 | $1.13 \times 10^{-3}$ |
| VWB3 | dip3 | 4.31 | 2.21 | $2.41 \times 10^{-4}$ |
| VWB3 | dip4 | 4.25 | 1.58 | $3.67 \times 10^{-4}$ |
| IT'IS3 | dip1 | 0.55 | 0.33 | $4.94 \times 10^{-5}$ |
| IT'IS3 | dip2 | 1.35 | 0.97 | $3.78 \times 10^{-4}$ |
| IT'IS3 | dip3 | 0.21 | 0.19 | $1.84 \times 10^{-5}$ |
| IT'IS3 | dip4 | 0.21 | 0.21 | $9.59 \times 10^{-6}$ |

(a) non-AMR means — Calibration

| Model | Dipole | Dist-mm | Angle-deg | Residual-Variance |
|---|---|---|---|---|
| SimNIBS3 | dip1 | 0.86 | 2.59 | $2.35 \times 10^{-4}$ |
| SimNIBS3 | dip2 | 0.83 | 2.95 | $4.66 \times 10^{-4}$ |
| SimNIBS3 | dip3 | 0.64 | 1.89 | $1.27 \times 10^{-4}$ |
| SimNIBS3 | dip4 | 1.06 | 0.82 | $3.92 \times 10^{-4}$ |
| VWB3 | dip1 | 0.78 | 2.38 | $2.10 \times 10^{-4}$ |
| VWB3 | dip2 | 0.84 | 2.92 | $4.33 \times 10^{-4}$ |
| VWB3 | dip3 | 0.59 | 1.72 | $1.10 \times 10^{-4}$ |
| VWB3 | dip4 | 0.97 | 0.74 | $3.25 \times 10^{-4}$ |
| IT'IS3 | dip1 | 0.18 | 0.26 | $1.77 \times 10^{-5}$ |
| IT'IS3 | dip2 | 0.41 | 0.76 | $3.24 \times 10^{-4}$ |
| IT'IS3 | dip3 | 0.08 | 0.14 | $1.18 \times 10^{-5}$ |
| IT'IS3 | dip4 | 0.10 | 0.10 | $4.31 \times 10^{-6}$ |

(b) non-AMR Standard Deviations — Calibration
